# SLAFIT: Viral fitness inference and structural variant detection from TGS data

**DOI:** 10.64898/2026.10.11.758307

**Authors:** Syed Muhammad Umer Abdullah, Dehan Cai, Jonathan Daniel Ip, Kelvin Kai-Wang To, Yanni Sun

**Affiliations:** Department of Electrical Engineering, City University of Hong Kong, Tat Chee Avenue, Kowloon, Hong Kong Special Administrative Region, China; State Key Laboratory of Emerging Infectious Diseases, Carol Yu Centre for Infection, Department of Microbiology, Li Ka Shing Faculty of Medicine, The University of Hong Kong, 102 Pokfulam Road, Hong Kong Special Administrative Region, China

## Abstract

Evolution is driven by genetic variation at multiple scales, from single-nucleotide variants (SNVs) to structural variants (SVs). Within populations, nonrandom associations among SNVs, quantified by linkage disequilibrium (LD), must be considered when inferring fitness from sequencing data. Characterizing SVs, SNVs, and the LD between SNVs therefore provides complementary insights into the genetic basis of evolutionary change. Although third-generation sequencing technologies facilitate the characterization of genomic variation, finite read lengths often limit the resolution of LD across entire genomes. Here, we present SLAFIT, a pipeline for linkage-aware fitness inference and SV detection from third-generation sequencing data. The pipeline reconstructs genome-wide haplotypes and their frequencies from third-generation sequencing reads to infer SNVs and estimate LD. When time-series data are available, it also infers fitness. An established SV detection tool is incorporated into the same workflow. Simulations support accurate LD and fitness inference under tested conditions, with reduced computation time through parallelization. Applications to enterovirus C104 and herpes simplex virus 1 reveal selection signals in predicted epitopes and recover a known inversion, respectively. Together, these results demonstrate the utility of SLAFIT for studying viral evolution through complementary analyses of fitness and structural variation.

## Introduction

Single nucleotide variants (SNVs) and structural variants (SVs) are fundamental sources of genomic evolution. SNVs alter individual genomic loci whereas SVs involve larger changes of at least 50bp in length^1^, including insertions, deletions, inversions, duplications and translocations. Both classes of variation can have major biological consequences. SNVs can alter phenotypes such as antimicrobial resistance^2^ in bacteria, and immune escape^3,4^ in viruses. SNVs can also alter essential biological functions, affect the structure of the organism, and modify other factors essential for survival, thereby directly affecting the replicative potential of an organism. SVs influence pathogenicity and antimicrobial resistance in pathogens^5,6^. SVs are also known to cause diseases, alter heritability, and affect environmental adaptation^7–9^. Together, these provide the raw material through which microbial and viral populations adapt to new hosts, drugs, immune pressures and ecological niches.

However, evolution is not driven only by individual SNVs acting in isolation. SNVs occur on genomic backgrounds, and their effects often depend on which other SNVs are present on which background. This non-random association between SNVs, known as linkage disequilibrium (LD), captures how SNVs are genetically linked within populations. Intuitively, LD quantifies the deviation of the observed pairwise occurrence of two SNVs from the pairwise occurrence expected under genomic independence. Biologically, LD reflects the history of mutation, recombination, selection, genetic drift, and population structure. LD can reveal whether SNVs are spreading together as part of the same successful lineage, whether neutral SNVs are hitchhiking with beneficial ones, or whether combinations of SNVs interact to increase or reduce fitness. Thus, LD provides information that cannot be obtained by considering SNVs independently.

LD is observed abundantly in nature and has been found to affect evolution in a number of human pathogens, including the acquisition of immune escape SNVs in influenza^10^ and the human immunodeficiency virus 1 (HIV-1)^11^. LD can also distort the observed prevalence of SNVs. LD between neutral and beneficial SNVs can cause the neutral SNVs to increase in prevalence while hitchhiking on the beneficial SNVs^12^. Conversely, beneficial SNVs can decrease each other’s prevalence because of LD^13^. Thus, the true fitness, or replicative potential, of SNVs in LD cannot be ascertained without quantifying the effect of LD on SNV prevalence^11^. LD, in addition to characterizing fitness, is required in the estimation of population divergence, calculation of population size, carrying out genome-wide association studies, and performing genomic prediction^14–17^.

High-throughput genomic sequencing, including next-generation sequencing (NGS)^19^ and third-generation sequencing (TGS), has played a transformative role in our ability to observe genomic evolution. TGS technologies, comprising Pacific Biosciences (PacBio) sequencing and Oxford Nanopore Technologies (ONT) sequencing, enable detailed observations of genomic evolution^20–22^. PacBio sequencing is known for its high accuracy which enables precise SV detection, targeted sequencing, accurate sequence assembly, and full-length transcriptome recovery^23–27^. ONT sequencing is popular for its large read lengths, low turnaround time, and low coverage requirement, which makes it ideal for use in onsite sequencing with limited time and resource availability^29–31^.

While TGS technologies can provide real-time, high resolution information about SNVs and SVs, the reads usually do not span complete viral genomes. As a result, the observable LD information is restricted to only those SNV pairs that are covered by the same read. The LD information remains unobservable for SNV pairs separated by distances exceeding the read length. Existing methods for LD inference from NGS data^36,37^ can, in principle, be adapted to work on TGS data, but these have slow computational speeds, model constraints, and scalability limitations, typically handling systems with no more than a few tens of loci. To the best of our knowledge, no method exists that takes TGS reads as input, detects both SNVs and SVs, and calculates the unobservable LD between SNVs. Consequently, LD information from the growing volume of TGS data remains only partially observed. Thus, there is a pressing need for a method that can estimate LD across the entire genome from TGS data in addition to detecting SNVs and SVs.

We present SLAFIT, a pipeline with the scope of SV detection, LD estimation, and fitness inference from TGS data. SLAFIT takes FASTQ files of TGS reads sampled at single or multiple evolutionary time-points as input; and outputs SNVs, SVs, and LD matrices at each time-point, together with fitness estimates when time-series data is available. SLAFIT makes no prior assumptions about the data and can identify the presence or absence of SVs, SNVs, and LD. In addition to unifying these analyses within a single framework, SLAFIT extends existing methodologies by enabling full-genome LD estimation from TGS reads and improving the efficiency of fitness inference. Because individual TGS reads provide partial information across a limited part of the genome, SLAFIT leverages a haplotype reconstruction based-approach. This method infers full-length haplotypes and their prevalence values that best explain the TGS reads, thereby enabling the calculation of LD across the entire genome. Furthermore, to accommodate high-throughput sequencing volumes, we developed a multiprocessing- and graphics processing unit (GPU)-based implementation of a fitness inference framework, MPL^11^, that reduces computational time compared to the original framework, particularly as genome size and number of sequences increase. We demonstrate the performance of SLAFIT for LD estimation on simulated data and show that fitness inference is robust across a range of read-sampling conditions. We also evaluate the false discovery rate of SV detection on simulated data and demonstrate that the method performs well.

To demonstrate the real-world utility of SLAFIT, we applied the pipeline to data from two viruses. We used SLAFIT for fitness inference on an enterovirus C104 (EV-C104) dataset. EV-C104 is a single-stranded ribonucleic acid (ssRNA) virus that rapidly accumulates SNVs via error-prone replication and recombination. This virus lacks an effective vaccine and can cause chronic infection in immunocompromised patients. We utilized longitudinal infection data from this virus and showed that

SLAFIT infers SNV fitness values that strongly align with computational predictions of immune escape. We also performed SV detection on data from the herpes simplex virus 1 (HSV-1). This virus is one of the most common human pathogens and does not have a vaccine. Identification of variation in the viral genome may lead to better understanding of its structure and vaccine development. We demonstrate that the proposed SV detection scheme is robust against reduction in the read-length, capturing structural variation profiles that match known baselines.

## Methods

### Overview

SLAFIT addresses two main problems, first being the detection of SVs from TGS reads. cuteSV^38^ was chosen for SV detection as it has been shown to consistently perform well over a range of test cases^39^. The second problem addressed is the reconstruction of unobservable LD values between SNVs that exist at distances greater than the read-length (referred to as inter-read LD values). To calculate the inter-read LD values, SLAFIT first uses RVHaplo^40^ to perform haplotype reconstruction. RVHaplo was selected as it has been shown to be sequencing platform-independent and has demonstrated good performance on real datasets of HIV-1 and HCV. RVHaplo takes in TGS reads and returns reconstructed haplotypes as well as the corresponding haplotype prevalence values, which are used to calculate the LD matrix. SNVs are calculated from the reconstructed haplotypes and returned with the LD estimate. When time-series data is available, the LD matrices are aggregated over all time-points via the proposed implementation of the MPL framework to obtain fitness values. A schematic of the pipeline is presented in Figure 1.

**Figure 1.**
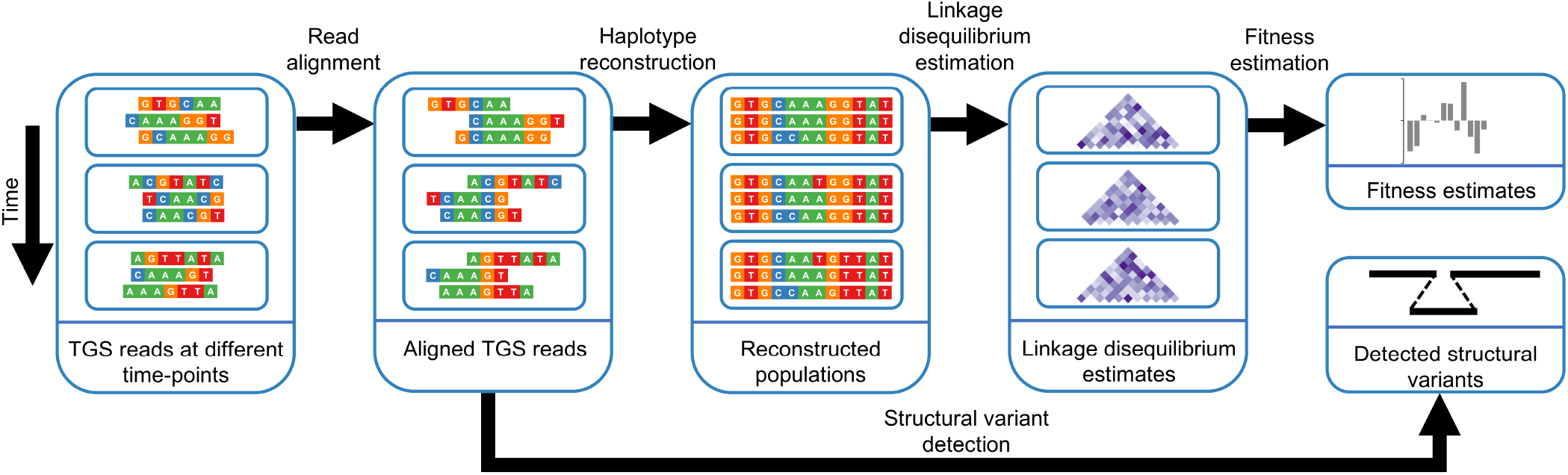
SLAFIT pipeline schematic. 1. TGS reads are obtained at each time-point from an evolutionary study. 2. TGS reads are aligned to a reference sequence to obtain Sequence Alignment Map (SAM) files at each time-point. 3. Haplotype reconstruction of the SAM files yields a set of full-length sequences and their corresponding prevalence values at each time-point. 4. LD matrices are obtained from the reconstructed sequences. 5. SV detection is performed via cuteSV and fitness estimates are obtained via the MPL equation.

### Read alignment and SV detection

The input to the pipeline are TGS reads sampled either at a single time-point, or at *K* + 1 time-points *{t*_0_, *t*_1_, …, *t*_*K−*1_, *t*_*K*_*}* in a time-series experiment. These reads are first aligned to a reference sequence, which is either user-supplied or obtained by taking the consensus of the reads sampled at the first time-point. Reference-based alignment of the input reads is performed via minimap2^41^. Subsequently, SV detection is performed on the aligned reads via cuteSV to obtain SVs captured by the reads.

### LD computation and fitness inference

The aligned TGS reads can contain spurious SNVs as a result of the high error rate of TGS. These reads are supplied to RVHaplo which first performs data-driven error correction to differentiate real SNVs from sequencing errors. The software subsequently performs haplotype reconstruction on the error-corrected reads and returns full-length reconstructed sequences and the prevalence values of the individual sequences. If the data contains a single dominant haplotype and other haplotypes occur at prevalence values that cannot be detected by RVHaplo, the software prompts the user that reconstruction cannot proceed. In this case the consensus sequence is used as a full-length reconstructed sequence as this is a good approximation for the actual population. The reconstructed sequences are then aligned against the reference sequence via Multiple Alignment using Fast Fourier Transform (MAFFT)^42^. The aligned, reconstructed sequences are supplied to the inference framework in which the entry of the LD matrix at loci *i* and *j, D*_*i j*_(*t*_*k*_) is computed as

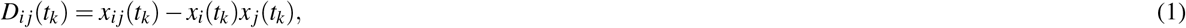

where *x*_*i*_(*t*_*k*_) and *x* _*j*_(*t*_*k*_) represent the single mutant allele frequencies at loci *i* and *j* respectively, *x*_*i j*_(*t*_*k*_) represents the double mutant allele frequency at loci (*i, j*), and *t*_*k*_ represents time-point *k*. The allele frequencies are obtained by counting the individual and pairwise occurrence of nucleotides that differ from the reference nucleotide at a particular locus. Fitness inference is subsequently performed via our proposed parallelized implementation of MPL. The estimated fitness values, single mutant allele frequencies, and the SNV counts are output by the pipeline.

### Parallel processing

Scalability can become a challenge during the analysis of large volumes of TGS data, for example, technologies such as PacBio High-Fidelity (HiFi) sequencing can generate over one million reads of lengths up to 10kbp^43^. Although these reads typically sequence the same genomic region covered by individual reads and are not affected by the limitation of unobservable inter-read LD information, the substantial redundancy in sampling introduces the challenge of scalability. Addressing this challenge is therefore essential for efficient estimation of observable LD from TGS data, independent of the unobservable inter-read LD estimation problem.

To improve scalability, we made two modifications to the MPL framework. First, the single- and double-mutant allele frequencies at each time-point are calculated on independent processor cores. Second, matrix inversion and multiplication in the MPL equation (Equation 1 in Sohail *et al*. 2021^11^) are carried out on a GPU, which is better suited to handle large-scale matrix operations. Together, these enhancements offer improved scalability compared to the original implementation, particularly as the number of sequences increases (Supplementary Figure S1).

### Overview of evaluation

We evaluated SLAFIT on both simulated and real data. The simulated data tests were designed to answer three main questions: (i) how robust SLAFIT is to degraded sequencing information, (ii) how well the method performs compared to existing approaches, and (iii) how well it scales to larger systems. The real data tests were designed to investigate whether SLAFIT was able to infer biologically-meaningful insights from the data. To assess robustness, we first used simulated data with known ground truth population composition and corresponding strain-level fitness values. In practice, these parameters are not observed directly and must be inferred from sequencing reads, which contain only partial and noisy information about the underlying genomes. We refer to this as information loss during read generation. This information loss arises because reads span only part of a genome, sequencing coverage may vary, and sequencing errors can corrupt the observed information. To systematically study the effect of information loss, we generated TGS reads under varying read lengths, coverage levels, and error rates, and then applied SLAFIT to infer strain-level fitness from the resulting reads. This allowed us to evaluate performance as the information available in the reads became increasingly degraded. Because the accuracy of fitness inference hinges on the recovery of LD, we also evaluated the accuracy of LD inference under these settings. We then compared SLAFIT against an NGS based scheme to quantify the gain in performance when using TGS reads. Furthermore, we performed comparative benchmarking to compare SLAFIT with a state-of-the-art linkage-aware fitness inference method. To assess scalability, we further examined performance as the genome length and number of sequences were increased. Finally, we applied SLAFIT to a real dataset to demonstrate its practical utility in generating biologically meaningful insights. We evaluated the SV detection component separately. First, using simulated data with no SVs, we estimated the false positive rate. We then tested SV detection on real data while varying TGS read length to simulate degraded input quality. These experiments showed that the method could detect a known SV even when the length of the SV was substantially longer than the maximum read length. We additionally identified putative SVs supported by multiple reads, which may warrant further biological validation.

### Simulated data

We generated simulated data with parameters guided by evolving viral populations. Data generation was performed via the Wright-Fisher model, which assumes non-overlapping populations of a finite number of individuals^44,45^. Unless otherwise specified, a population size of *N* = 1000 individuals of length *L* = 4500bp was chosen. At the first time point of evolution, *N* copies of a random sequence of length *L* were generated. The base ‘A’ was considered the wild type (WT) and assigned 0, whereas the bases {C,G,T} were considered mutants and assigned 1. The bi-allelic model used here is for the sake of simplicity and can be readily expanded to a 4-allelic model (Sohail *et al*. 2021^11^ p. 481). The generated population was cloned and mutated based on the Wright-Fisher evolutionary dynamics. Every cloning step corresponded to one generation. At each generation, selective pressure was simulated on predefined loci to choose for fitter SNVs. This evolution was simulated for 550 generations, from which *n* = 10 individuals were sampled every Δ*t* = 50 generations, starting at the 51st generation. The subsampling of sequences and time emulates real sequencing, in which a sample cannot represent the entire population, and the samples cannot be taken at every generation. For every sequence represented by a binary string 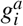, the sequence fitness *f*_*a*_ was assumed to follow an additive model, given by

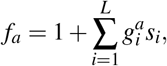

where *s*_*i*_ represents the selection coefficient of the mutant allele at locus *i*. The additive fitness model is a simplification that assumes no epistasis between loci. The MPL equation is based on the additive fitness model, and we use the model for consistency. The SNV probabilities from WT to mutant (*µ*_WT,mut_) and mutant to WT (*µ*_mut,WT_) were assumed to be equal and assigned a value *µ* = 10^*−*4^ per base per generation. The number of beneficial, deleterious, and neutral SNVs were {180, 180, 4140}. The beneficial and deleterious SNVs were assigned the same magnitude of selection coefficients, 0.025, with opposite signs, while the neutral SNVs were assigned zero-valued selection coefficients. The beneficial and deleterious SNVs were uniformly distributed across the length of the genome. We generated three datasets with genome length {9000, 13500, 18000}, and three datasets with number of sequences {100, 1000, 10000}. For datasets with genome length {9000, 13500, 18000}, the number of beneficial, deleterious, and neutral SNVs were {360, 360, 8280*}, {*540, 540, 12420}, and {720, 720, 16560}, respectively, which were distributed uniformly across the genome. All datasets were generated without SVs. 100 Monte Carlo runs were generated for each dataset to establish statistical confidence in the results.

TGS was simulated for the error profile of the Oxford Nanopore R10.4.1 system via badread^46^. For each time-point, reads of mean length 1500 bp were generated with a coverage of 50X, and mean/maximum identity 90*/*100 (Supplementary Figure S2). The term “coverage” is used to refer to the population-level coverage, given as *C* = *lR/L*, where *C* =coverage, *R* =number of reads, *l* = read length, and *L* = genome length. The parameters “junk reads”, “random reads”, “chimeras”, and “glitches” were set to 0, and no adapter sequence was used on either end. These parameters correspond to a fairly clean sequencing run. For testing robustness to read parameters, we generated additional datasets with different parameters corresponding to sequencing runs with relatively degraded quality. Two datasets were generated with read length {500, 1000}, two datasets with coverage {25X, 35X}, and two datasets were generated with mean identity {85, 80}.

For inference, we set the value of the regularization parameter *γ* in the MPL framework to 20. This parameter is heuristic and inference has been shown to be stable for a range of its values (Supplementary Fig. S2 in Abdullah *et al*. 2026^47^). Estimation was performed on only those loci on which the single mutant allele frequency exhibited at least 10% variation, which is the least count when sampling *n* = 10 individuals. The single mutant allele frequencies were computed from the simulated population data.

### EV-C104 data

We performed SNV-level fitness inference on a real dataset of prolonged EV-C104 infection^48^ to demonstrate the ability of SLAFIT to infer biologically meaningful insights. The clinical timeline of the infection is presented in Supplementary Figure S3. Briefly, the dataset was obtained over a course of 216 days from a patient with multiple myeloma relapse. The data consists of 57899 *±* 38364 ONT MinION reads of length 889 *±* 107 bp with a coverage of 7027X*±*5137X. The read-length used is much shorter than the genome-length (7390 bp) which means that much of the long-range LD information is unobservable (Supplementary Figure S4). Following Cassidy *et al*. 2022^48^, the consensus sequence of the first time-point (GenBank accession no. MZ092702) was used as reference and reads for all time-points were aligned against it as explained in the Methods section. Single mutant allele frequency trajectories, obtained via LoFreq^49^ indicated multiple co-occurring SNVs, which suggested the existence of LD in the data (Figure S5). Haplotype reconstruction was performed to obtain full-length sequences of the coding region (703 *−* 7323 bp) of the genome and selection coefficients obtained via a bi-allelic (WT/mutant) model. The choice of a bi-allelic model was made because the entropy values of the single mutant allele frequencies indicated that this model accurately captured most of the variation in the data (Supplementary Figure S6). Selection coefficient estimation was performed via the proposed implementation of MPL with the regularization parameter *γ* = 20. SNV probability of 2.97 *×* 10^*−*3^ per year estimated by Cassidy *et al*. 2022^48^ was used for WT-mutant transitions and three times this probability was used for the mutant-WT transitions to account for transitions from three possible mutants to the WT. Inference was performed on those loci where the single mutant allele frequency exhibited at least 1% variation. This threshold was chosen based on the sensitivity of RVHaplo, which has been demonstrated to accurately recover variants with as little as 1% prevalence.

### HSV-1 data

We used a real dataset of HSV-1^50^ to test the SV detection scheme. The dataset comprised 104524 ONT sequencing reads with mean length 1294 *±* 4100bp. 15 reads were ultra long, spanning more than 140kbp. The reads were aligned against the HSV-1 reference sequence (NCBI accession no. KU310657.1) and SV detection performed as described before.

## Results

### Experiment 1: LD estimation and fitness inference via SLAFIT is robust to varying sequencing parameters

In real-life sequencing scenarios, control over sequencing parameters is limited by time and resource constraints. For archival samples, analysis must rely on already sequenced datasets without any control over the sequencing parameters. We tested the performance of the fitness inference scheme of SLAFIT for various values of sequencing parameters to check the robustness to varying levels of noise in the data. We performed inference on the sequences of the ground truth population to use as a baseline (Figure 2**A**), and varied the read length (Figure 2**B** and Supplementary Figure S7**A**), coverage (Figure 2**C** and Supplementary Figure S7**B**), and error rate (Figure 2**D** and Supplementary Figure S7**C**). SLAFIT performed well for some degradation in the read quality. The performance demonstrated significant degradation when the error rate was set to 20% or a read length of 500bp was used. This is because errors in the reads or the information lost due to the smaller read length propagate into the reconstructed haplotypes and their prevalence values, which affects the estimated LD and the selection coefficients. Though these values of error rate and read-length are used for benchmarking purposes and may not be encountered when using modern sequencing machines. These results show that accuracy of fitness inference directly correlated with the data quality and can serve as a guideline for designing sequencing experiments.

**Figure 2.**
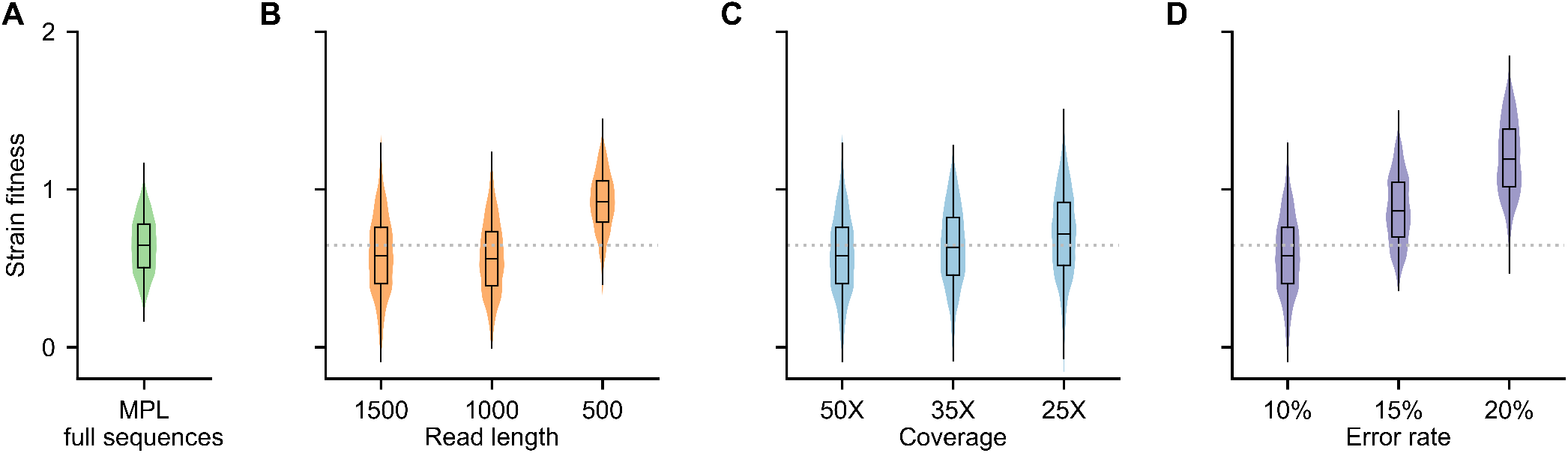
SLAFIT can perform inference robustly over a range of sequencing parameters. (A) A distribution of the strain-level fitness obtained from the ground truth population is shown. The strain-level fitness, estimated via SLAFIT is shown while varying the (B) read length, (C) sequencing coverage, and (D) error rate. The dotted gray line represents the median of the strain-level fitness of the ground truth population. Results are shown for 100 Monte Carlo runs. Each Monte Carlo run consisted of evolving populations of *N* = 1000 individuals of genome length *L* = 4500 bp. The genomes consisted of bi-allelic (WT and mutant) loci with equal forward and backward SNV probabilities set to *µ* = 10^*−*4^ per locus per generation. Alleles at 180*/*180*/*4140 loci were respectively beneficial/deleterious/neutral with selection coefficients +0.025*/ −* 0.025*/*0. The beneficial and deleterious mutants were uniformly distributed across the entire genome. One founder sequence was used to generate each population, which was allowed to evolve for 550 generations and sampled at Δ*t* = 50 generations. Unless otherwise stated, TGS reads of mean length 1500 bp were simulated with 10% error rate and 50X coverage.

In all the tested datasets, SNVs were spread across the genome, and long-range LD values were not directly observable via TGS reads. Nevertheless, the haplotype reconstruction scheme of the pipeline accurately reconstructed the full-length sequences, from which the unobservable long-range LD values were faithfully recovered, enabling accurate fitness inference. Supplementary Figure S8 shows Circos plots^51,52^ for LD values aggregated over the entire evolutionary history. Supplementary Figure S8 **A** and **B** demonstrate that the ground truth and inferred aggregated LD are similar whereas the inferred selection coefficients have the same sign. Supplementary Figure S8 **C** presents the summary statistics of the comparison of the inferred and ground truth aggregated LD over 100 Monte Carlo runs. We also quantified the accuracy of LD recovery via the Pearson correlation coefficient between the LD values obtained from the reconstructed sequences and the LD values obtained from the sequences of the ground truth population (Supplementary Figure S9). The recovered LD values exhibited strong correlation with the ground truth LD values. The relatively weaker correlation observed at the initial generations is because of lower variation in the data which affects haplotype reconstruction. The correlation values of the first two time-points are not shown as haplotype reconstruction failed to return haplotypes for most Monte Carlo runs due to low variation, and consequently, the consensus sequences of the reads were used in subsequent analysis of those time-points. The lack of variation at initial time-points is characteristic of natural infections in which the infecting strain initially dominates the population. For these time-points, other strains are either not present or present at very low prevalence values and consequently, the consensus strains can approximate the population well. For subsequent time-points, enough variation exists for population reconstruction to identify unique haplotypes.

### Experiment 2: Longer read length of TGS reads yields improved fitness estimates compared to NGS reads

We compared strain-level fitness estimates reported by TGS and NGS to demonstrate the advantage of the longer reads of TGS. We obtained TGS reads and performed haplotype reconstruction as described in the Methods section. We then used ART^53^ to obtain NGS reads with a coverage comparable to that of the TGS data. We simulated 14500 single-end NGS reads of 150bp using the error profile of the Illumina HiSeq 2500 platform and applied MPL-R^47^ to obtain fitness estimates.

Figure 3 demonstrates the distribution of the strain-level fitness estimates obtained via SLAFIT and MPL-R. SLAFIT yielded estimates that more closely matched those obtained from the ground truth population as compared to MPL-R. The Pearson correlation coefficients of the respective estimates also indicated that SLAFIT yielded fitness values that were more strongly associated with the ground truth as compared to MPL-R (Supplementary Figure S10).

**Figure 3.**
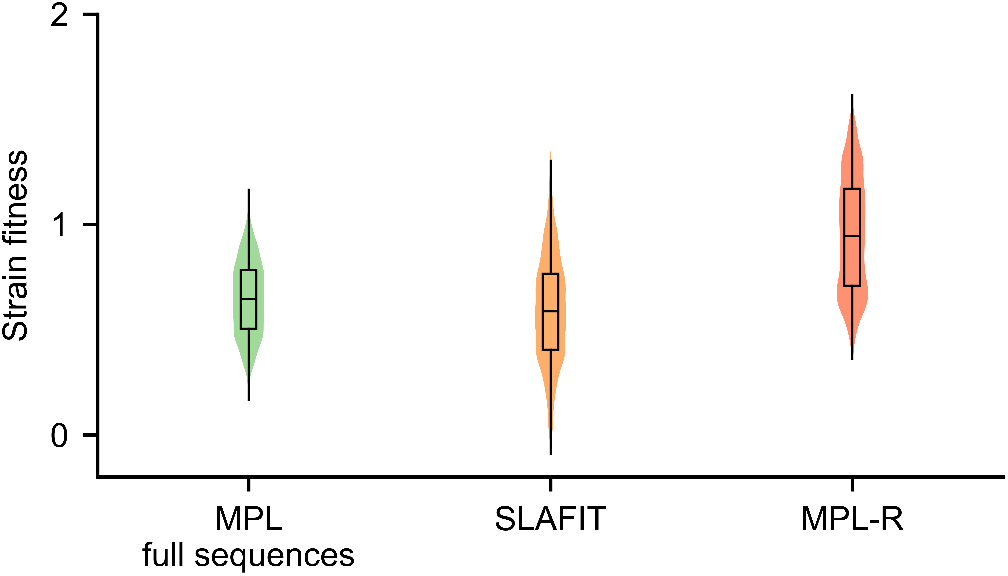
SLAFIT is able to reconstruct fitness values from TGS reads more accurately as compared to MPL-R, an NGS-based scheme. The distributions are shown for 100 Monte Carlo runs. Each Monte Carlo run consisted of evolving populations of *N* = 1000 individuals of genome length *L* = 4500 bp. The genomes consisted of bi-allelic (WT and mutant) loci with equal forward and backward SNV probabilities set to *µ* = 10^*−*4^ per locus per generation. Alleles at 180*/*180*/*4140 loci were respectively beneficial/deleterious/neutral with selection coefficients +0.025*/ −* 0.025*/*0. The beneficial and deleterious mutants were uniformly distributed across the entire genome. One founder sequence was used to generate each population, which was allowed to evolve for 550 generations and sampled at Δ*t* = 50 generations. TGS reads of mean length 1500 bp were simulated with 10% error rate and 50X coverage. 14500 single-end NGS reads of 150bp were simulated to obtain similar coverage as the TGS reads.

To ensure a fair comparison, we attempted to use the same haplotype reconstruction software for both TGS and NGS data, but RVHaplo performed poorly on NGS data, and Quasirecomb^54^, the software used in MPL-R, was too computationally expensive to run on TGS data. This is expected as software designed for either technology is optimized to handle the data of only its intended technology. Therefore we used each reconstruction software for the specific technology for which it was developed. While the usage of different software for haplotype reconstruction introduces a confounder, the difference in performance between the two technologies is qualitatively similar to what is observed when TGS reads of different length are used, which indicates that the read length is likely the major factor for the difference in performance between the data from the two technologies. This can be explained in terms of the error rates of both technologies as well. NGS typically has a lower error rate compared to TGS, so for the same read length, NGS is expected to yield better performance compared to TGS. Hence the observed difference in performance is likely because of TGS capturing more information from the reads due to its greater read length.

### Experiment 3: SLAFIT outperforms existing LD-aware fitness inference

Inference methods that take into account LD are designed only for NGS data with a few SNVs and scale poorly when applied to larger systems^37,55,56^. These methods have been shown to require several hours to run on a single Monte Carlo run^47,57^ and were not used for comparison. The only LD-aware method that can process sequences of length 4500bp without requiring a large wall clock time is the method of Li and Barton 2023^58^ (subsequently referred to as LB). We used single mutant allele frequencies obtained from the TGS reads as input to LB, but it performed poorly compared to SLAFIT (Figure 4). Thus to our knowledge, SLAFIT remains the only method that can efficiently perform accurate LD-aware fitness inference.

**Figure 4.**
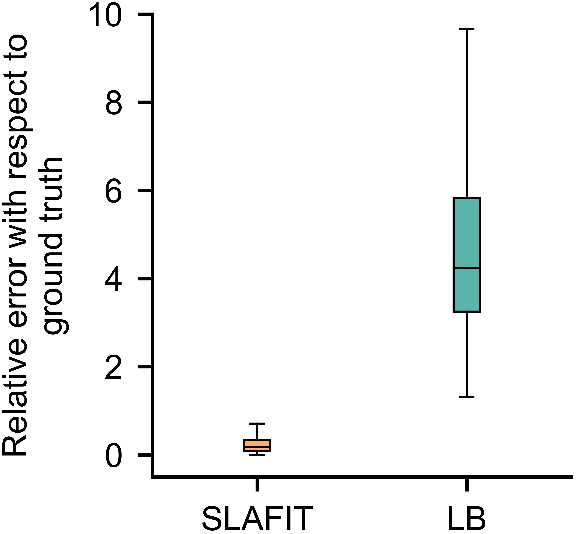
Fitness values inferred via SLAFIT have lower relative error with respect to the ground truth as compared to the method of Li and Barton 2023^58^, the only LD-aware inference method that can scale up to systems of kpb scale. The relative error is defined as the absolute difference between the strain fitness inferred by the method under test and MPL on the ground truth sequences divided by the absolute value of the fitness inferred by MPL on the ground truth sequences. Results are shown for 100 Monte Carlo runs. Each Monte Carlo run consisted of evolving populations of *N* = 1000 individuals of genome length *L* = 4500 bp. The genomes consisted of bi-allelic (WT and mutant) loci with equal forward and backward SNV probabilities set to *µ* = 10^*−*4^ per locus per generation. Alleles at 180*/*180*/*4140 loci were respectively beneficial/deleterious/neutral with selection coefficients +0.025*/ −* 0.025*/*0. The beneficial and deleterious mutants were uniformly distributed across the entire genome. One founder sequence was used to generate each population, which was allowed to evolve for 550 generations and sampled at Δ*t* = 50 generations. TGS reads of mean length 1500 bp were simulated with 10% error rate and 50X coverage.

### Experiment 4: SLAFIT reveals insights from prolonged infection data consistent with biological knowledge

We applied SLAFIT on prolonged intrahost EV-C104 infection data. Haplotype reconstruction of the data returned {15, 13, 8, 11} unique haplotypes at each of the time-points respectively. We analyzed the data via SLAFIT and as a benchmark, also performed LD-independent inference via SLAFIT by setting the off-diagonal entries of the LD matrix to zero. A plot of the ratio of the selection coefficients for nonsynonymous SNVs inferred via SLAFIT and SLAFIT (LD-independent) indicates that SLAFIT reports much higher selective pressure on five SNVs as compared to SLAFIT (LD-independent) (Figure 5). Experimental evidence of the significance of these SNVs is unavailable and the virus has no known lab-validated epitopes. Therefore, to validate our findings, we predicted the B-cell epitopes of the virus via the Immune Epitope Database (IEDB)^59^ using the Bepipred Linear Epitope Prediction 2.0 tool^60^ with default threshold of 0.5. Four of the five SNVs classified as beneficial by SLAFIT and neutral by SLAFIT (LD-independent) were observed to occur in the predicted B-cell epitopes (Supplementary table S1). This is consistent with biological knowledge that epitopes are under immune pressure and nonsynonymous SNVs in epitopes have high fitness as these likely assist in immune escape. This demonstrates that SLAFIT, via the recovery of unobservable LD information, can infer insights from real data that are missed if LD is not considered. Here we must clarify that the epitopes were predicted computationally due to the unavailability of experimentally-determined epitopes of the virus. These predictions were generated to identify whether the nonsynonymous SNVs lie in regions predicted to be surface-exposed and immunogenic. However, these predictions represent computational concordance, not experimental validation. We therefore use the results of these predictions as a biologically informed benchmark to assess the validity of our results, rather than as concrete evidence of immune escape. Future work employing neutralization assays can experimentally validate the significance of these SNVs.

**Figure 5.**
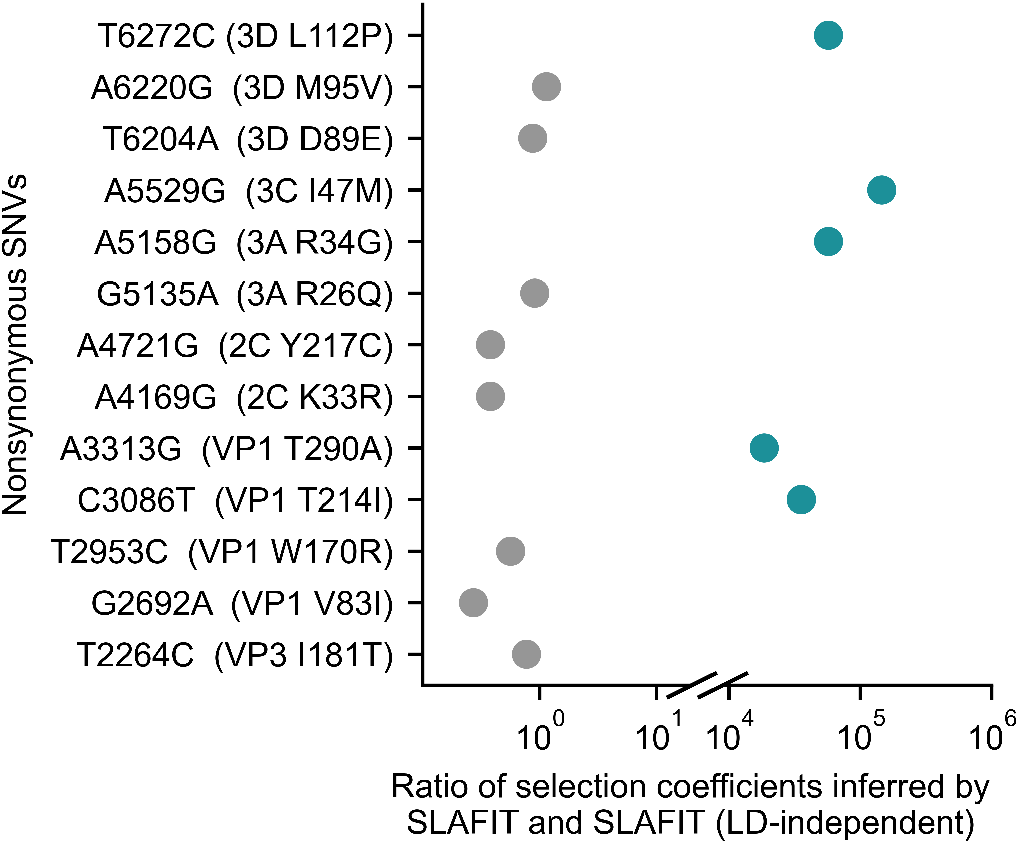
LD-aware fitness inference via SLAFIT reports biologically concordant insights. SLAFIT infers a number of nonsynonymous SNVs as non-neutral (blue) which are identified as neutral by SLAFIT(LI). Four of these SNVs map to B-cell epitopes predicted via IEDB and are described in Supplementary table S1.

The LD values obtained via SLAFIT, aggregated over the complete evolutionary history, were mostly negative, which suggested clonal interference between nonsynonymous SNVs (Supplementary Figure S11). This observation suggested competition between strains within the population which was also evident by the decreasing number of reconstructed haplotypes with time. The decreasing single mutant allele frequencies (Supplementary Figure S5) and the decreasing haplotype diversity (Supplementary Figure S12) also indicated strains being outcompeted and removed from the population.

### Experiment 5: SV detection via cuteSV is robust and consistent with known results

We performed SV detection on the dataset of HSV-1 by first considering all available reads. The SV in the U_L_ region reported in Saranathan *et. al* (2022) was not detected, which is expected, since the number of reads that can cover the entire length of the U_L_ region (110026 bp) was 19 (Supplementary Figure S13). We were able to detect SVs in the U_S_ region, out of which two SVs matched the inversion in the U_S_ region reported in the original study. This dataset, however, had ultralong reads (exceeding the length of the U_S_ region), which are not always available in TGS datasets. To validate the robustness of SV detection to more realistic reads, we sequentially removed reads greater than a fraction (0.75, 0.50, 0.25, and 0.125) of the length of the U_S_ region and repeatedly performed SV detection. The inversion in the U_S_ region was detected even when the maximum length of the reads was as low as 0.25 that of the region. This shows that the detected SVs are consistent with the SV reported in the original study. The SVs have been summarized in Table 1.

**Table 1.** SVs detected from HSV-1 data under varying maximum read-length thresholds. The U_*S*_ region (reference) is shown for comparison.

| Max read length (bp) | SV detected | Start | End | Read support |
| --- | --- | --- | --- | --- |
| NA (reference) | $U_S$ region | 137276 | 150359 | NA |
| 184240 | SV 1 | 135457 | 150377 | 14 |
| 184240 | SV 2 | 137200 | 150373 | 11 |
| 9813 | SV 1 | 135457 | 150377 | 14 |
| 6542 | SV 1 | 137200 | 150373 | 11 |
| 3271 | SV 1 | 137200 | 150373 | 11 |
| 1635 | SV 1 | 137197 | 150376 | 10 |

## Discussion

In this work, we have presented SLAFIT for the detection of SVs and computation of fitness from TGS data. We have demonstrated the performance of SLAFIT on simulated data under varying read lengths, coverage values, and error rates. We have shown that fitness inference via SLAFIT is accurate, completes in a smaller wall clock time compared to MPL, and outperforms MPL-R, and LB, the only other LD-aware methods capable of scaling up to sequence lengths in the kbp range. SLAFIT yielded biologically consistent insights from EV-C104 infection data, detecting four strong nonsynonymous SNVs in computationally-predicted B-cell epitopes that were missed by an LD-independent approach. The SV detection scheme used in SLAFIT, cuteSV, detected SVs in an HSV-1 dataset even when the maximum read length was as short as one-quarter of the length of the target region. These results demonstrate that SLAFIT enables evolutionary analysis from TGS data that is not possible with other methods.

In terms of limitations, large genome lengths may require careful handling. The largest genome we tested SLAFIT on was 18000 bp in length. Based on the trend in performance, we expect the timing performance of SLAFIT to scale well for larger genomes, however, care should be taken to ensure all matrices are within the limits dictated by the system memory. SLAFIT is also limited by the usage of an additive fitness model. Fitness may not always be additive in nature and epistatic interactions between SNVs warrant the incorporation of triple- and quadruple-mutant allele frequencies. While mathematical frameworks exist for the incorporation of epistasis^61,62^, efficiently computing and storing quadruple-mutant allele frequencies for high-throughput data of systems of the scale of kbp remains a challenge. Further research can attempt to address this limitation. Another limitation is the use of the 1% threshold for differentiating between true variants and sequencing errors. While this threshold is consistent with published benchmarks on RVHaplo, sequencing conditions and strain abundance may vary between datasets. To obtain an accurate threshold for a particular dataset under a particular set of sequencing conditions, simulations may need to be carried out to ascertain the exact prevalence threshold below which it becomes challenging to distinguish actual variants from errors. Future work can investigate this question.

One of the most valuable tools available to clinicians is whole-genome third-generation sequencing of time-series patient samples^35,63–67^. Commonly used pipelines are limited to consensus-sequence based analysis and cannot resolve the interaction between SNVs at the strain level. SLAFIT allows an integrated pipeline to obtain accurate insights from TGS data at a finer scale. The core of fitness inference of SLAFIT is RVHaplo, which enables the pipeline to be able to run on a number of sequencing machines. Fitness inference via SLAFIT is also designed to be modular, so users can swap individual software components with more familiar ones. The SV detection scheme of SLAFIT is based on a robust software, cuteSV, whose performance has been experimentally validated on multiple real datasets. By enabling strain-level evolutionary inference, SLAFIT enables more precise monitoring of pathogen evolution in real-world settings.

## Acknowledgements

This work was carried out on the High Performance Computing Cluster provided by the Department of Electrical Engineering and CityU HK Burgundy High Performance Computing system managed and provided by the Computing Services Centre at City University of Hong Kong. The authors would like to express gratitude to Prof. Runsheng Li, Prof. Ka Chun Wong, Prof. Xiangjun Du, Dr. Muhammad Ashraf Abdelhamid Abdelgawad, Dr. Abdurrashid Ibrahim Sanka, and Dr. Muhammad Saqib Sohail for their insightful comments that improved the content of this study.

## Data availability

Data for the EV-C104 and HSV-1 studies can be accessed from the repositories mentioned in the respective publications. Simulated data for testing SLAFIT can be accessed on the GitHub repository.

## Software availability

The code for SLAFIT is available with example data on GitHub at https://github.com/SMUAbdullah/SLAFIT.

