## supplementary_information for "SLAFIT: Viral fitness inference and structural variant detection from TGS data"

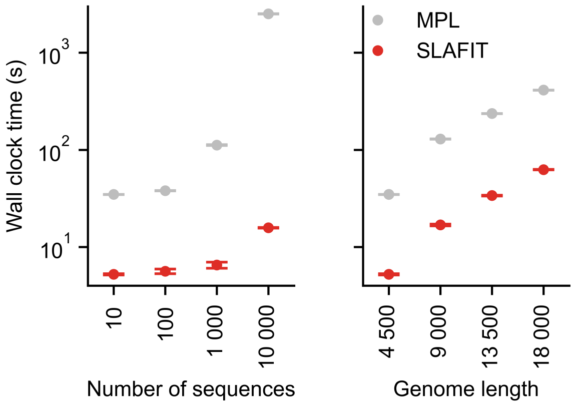


**Figure S1. Parallelized implementation of MPL supplied with SLAFIT requires lower wall clock time compared to the sequential implementation.** The wall clock time required by both implementations of MPL increases with the number of sequences (*left panel*) and genome length (*right panel*). The implementation used in SLAFIT consistently takes less time. Circles indicate the mean whereas the error bars indicate the standard error of the mean. Results are shown for 100 Monte Carlo runs. For the tests involving genome length, each Monte Carlo run consisted of evolving populations of $N = 1\,000$ individuals. For the tests involving number of sequences, the genome length was fixed at $L = 4\,500$ bp. The genomes consisted of bi-allelic (WT and mutant) loci with equal forward and backward SNV probabilities set to $\mu = 10^{-4}$ per locus per generation. For genome lengths $4\,500, 9\,000, 13\,500, 18\,000$ alleles at $180/180/4140, 360, 360, 8\,280, 540, 540, 12\,420$, and $720, 720, 16\,560$ loci were respectively beneficial/deleterious/neutral with selection coefficients $+0.025/-0.025/0$. The beneficial and deleterious mutants were uniformly distributed across the entire genome. One founder sequence was used to generate each population, which was allowed to evolve for 550 generations and sampled at $\Delta t = 50$ generations. TGS reads of mean length 1 500 bp were simulated with 10% error rate and 50X coverage. All tests were carried out on an NVIDIA RTX 4080 GPU on a system with an Intel Xeon 3.5 GHz CPU with 512GB memory.

**Table S1.** Nonsynonymous SNVs identified as highly fit via linkage-aware inference occur in computationally predicted B-cell epitopes.

| NT mutation | AA mutation | Gene | Predicted B-cell epitope | Gene function |
|---|---|---|---|---|
| C3086T | T214I | VP1 | SVVPLNTVDAGASNKY | Receptor-binding domain |
| A3313G | T290A | VP1 | RAADYVGRSVDYNQNSPVMIATAD | Receptor-binding domain |
| A5158G | R34G | 3A | VDSEEVREY | Viral replication |
| T6272C | L112P | 3D | TDGLEALDLT | Viral replication |

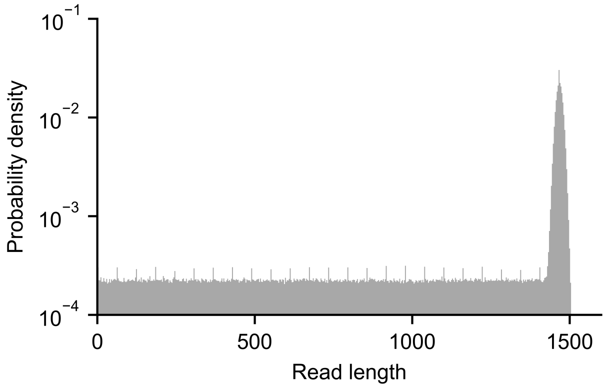


**Figure S2. Simulated TGS reads are generated with various lengths and provide different levels of information.** Read lengths are shown for 100 Monte Carlo runs. Each Monte Carlo run consisted of evolving populations of $N = 1\,000$ individuals of genome length $L = 4\,500$ bp. The genomes consisted of bi-allelic (WT and mutant) loci with equal forward and backward SNV probabilities set to $\mu = 10^{-4}$ per locus per generation. Alleles at $180/180/4140$ loci were respectively beneficial/deleterious/neutral with selection coefficients $+0.025/-0.025/0$. The beneficial and deleterious mutants were uniformly distributed across the entire genome. One founder sequence was used to generate each population, which was allowed to evolve for 550 generations and sampled at $\Delta t = 50$ generations. TGS reads of mean length 1 500 bp were simulated with 10% error rate and 50X coverage.

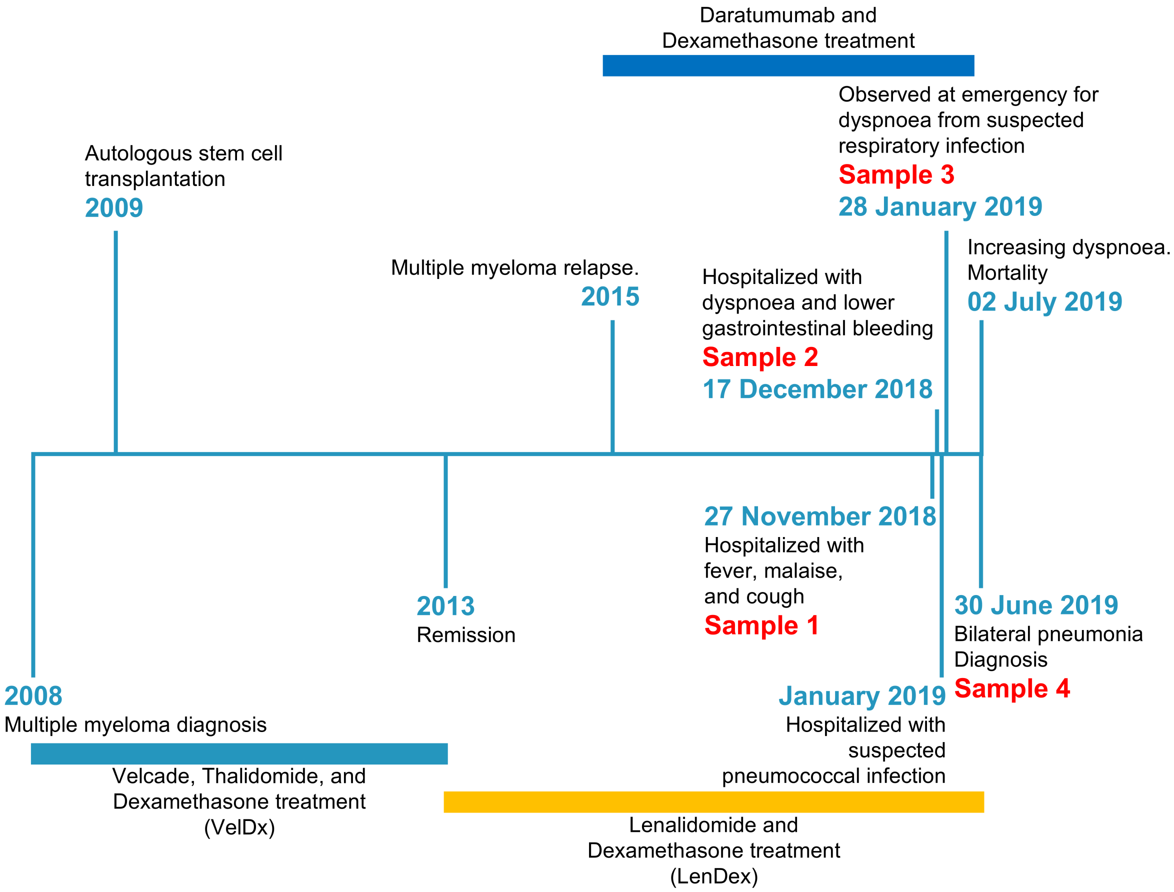


**Figure S3.** Clinical timeline of EV-C104 infection data.

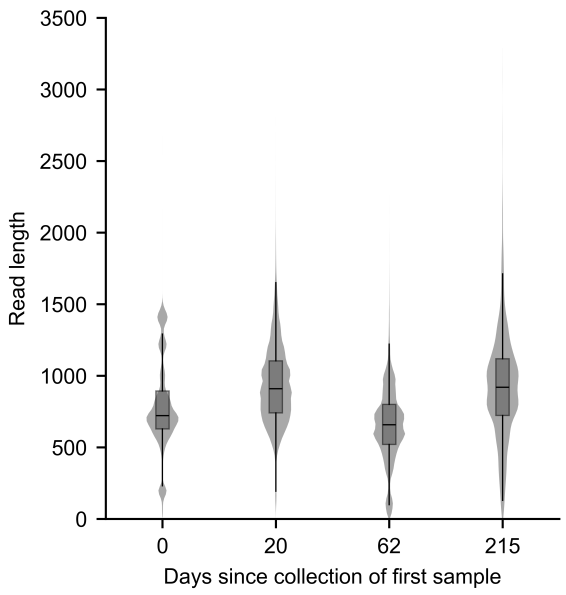


**Figure S4. Read-length used in the EV-C104 infection data is much shorter than the genome length.** TGS reads used to sample the EV-C104 infection data left much of the long-range LD information unobservable due to much shorter lengths compared to the genome-length.

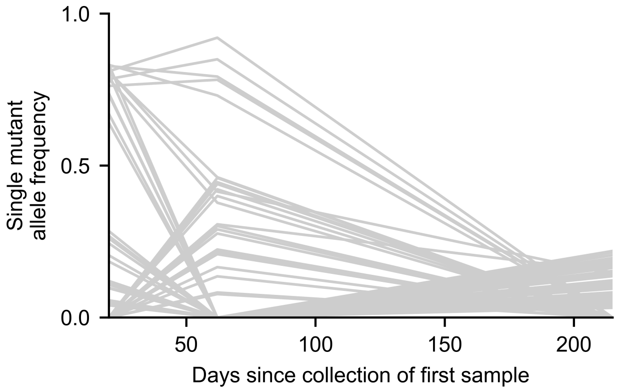


**Figure S5. EV-C104 infection data indicates the presence of LD.** Multiple co-occurring SNVs are observed in the EV-C104 infection data which suggests towards the existence of LD.

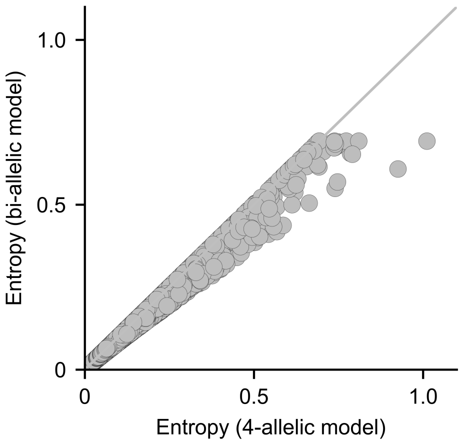


**Figure S6. Bi-allelic model for the EV-C104 infection data shows good association with the 4-allelic model.** Entropy values of the single mutant allele frequencies computed via bi- and 4-allelic models are strongly correlated.

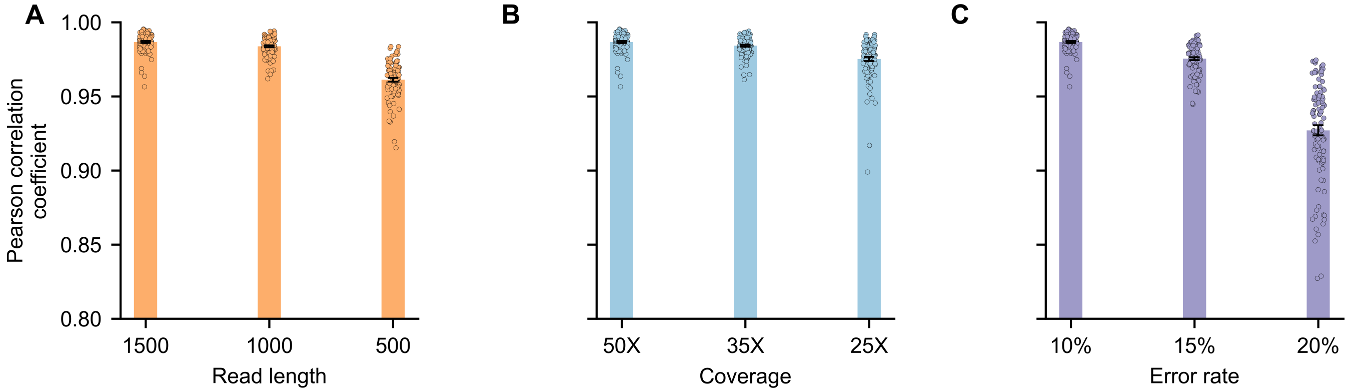


**Figure S7. Fitness inference via SLAFIT is robust over a range of sequencing parameters.** The strain-level fitness inference of SLAFIT is strongly associated with the ground-truth with varying (A) read length, (B) sequencing coverage, and (C) error rate. The bars show the mean Pearson correlation coefficient over 100 Monte Carlo runs. Each Monte Carlo run consisted of evolving populations of $N = 1\,000$ individuals of genome length $L = 4\,500$ bp. The genomes consisted of bi-allelic (WT and mutant) loci with equal forward and backward SNV probabilities set to $\mu = 10^{-4}$ per locus per generation. Alleles at $180/180/4140$ loci were respectively beneficial/deleterious/neutral with selection coefficients $+0.025/-0.025/0$. The beneficial and deleterious mutants were uniformly distributed across the entire genome. One founder sequence was used to generate each population, which was allowed to evolve for 550 generations and sampled at $\Delta t = 50$ generations. Unless otherwise stated, TGS reads of mean length 1 500 bp were simulated with 10% error rate and 50X coverage. The error bars show the standard error of the mean.

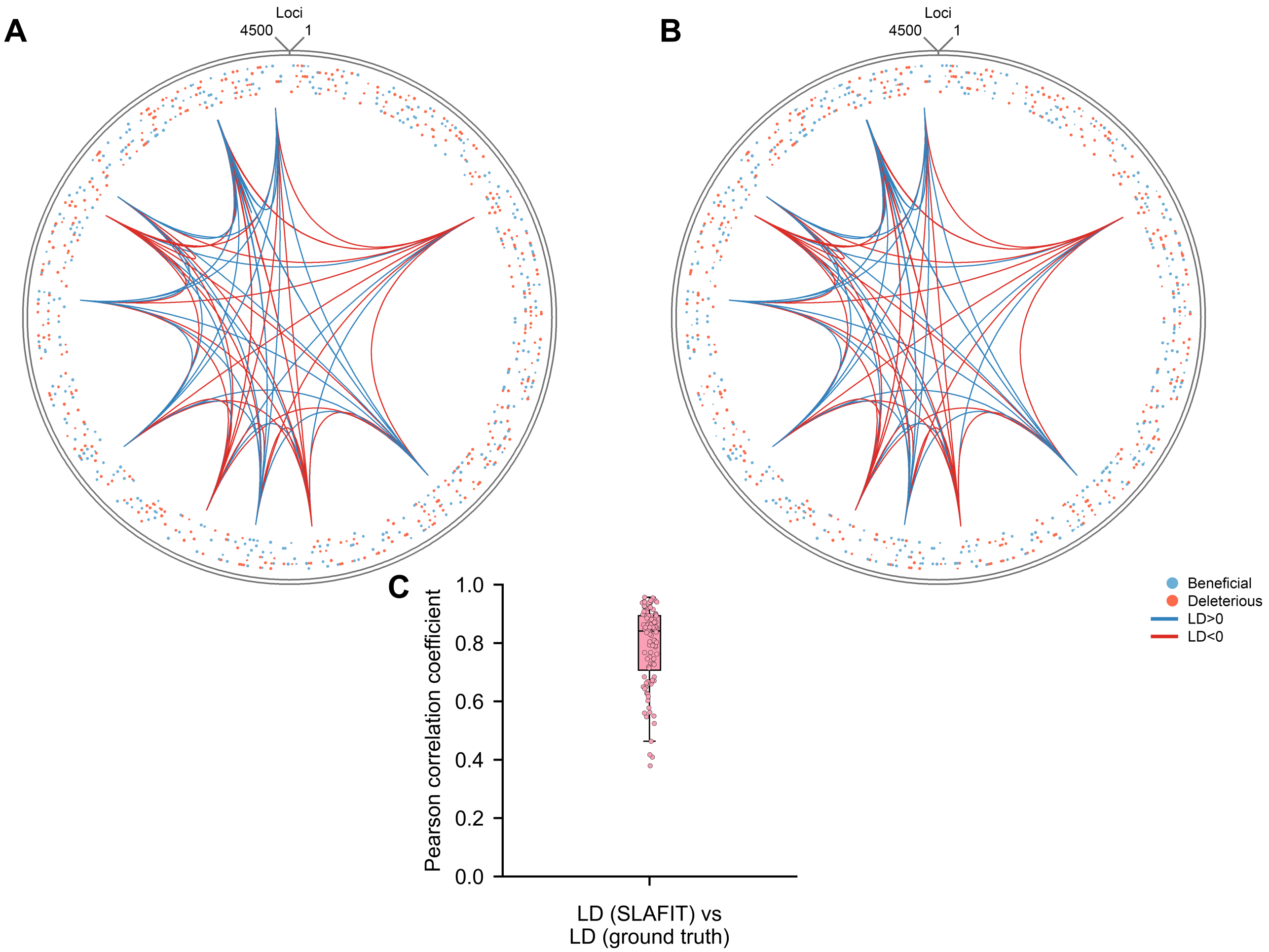


**Figure S8. SLAFIT accurately recovers long-range LD values.** (A) Aggregated LD values obtained from the ground truth population and selection coefficients inferred from those. (B) Aggregated LD values obtained via SLAFIT and selection coefficients inferred from those. (C) Aggregated LD values inferred via SLAFIT show good correlation with the ground truth aggregated LD values. Results in (A) and (B) are shown for a typical Monte Carlo run. log magnitudes of the inferred selection coefficients are indicated as the radii of the circles. Blue circles represent positive values, whereas red circles represent negative ones. Links between loci indicate LD values aggregated over all time-points. The widths of the link indicate the magnitude of the aggregated LD, and blue/red colors indicate positive/negative sign of the aggregated LD. Only the strongest 65 aggregated LD values are shown to avoid clutter. Results in (C) are shown for 100 Monte Carlo runs. Each Monte Carlo run consisted of evolving populations of $N = 1\,000$ individuals of genome length $L = 4\,500$ bp. The genomes consisted of bi-allelic (WT and mutant) loci with equal forward and backward SNV probabilities set to $\mu = 10^{-4}$ per locus per generation. Alleles at $180/180/4140$ loci were respectively beneficial/deleterious/neutral with selection coefficients $+0.025/-0.025/0$. The beneficial and deleterious mutants were uniformly distributed across the entire genome. One founder sequence was used to generate each population, which was allowed to evolve for 550 generations and sampled at $\Delta t = 50$ generations. Unless otherwise stated, TGS reads of mean length $1\,500$ bp were simulated with 10% error rate and 50X coverage.

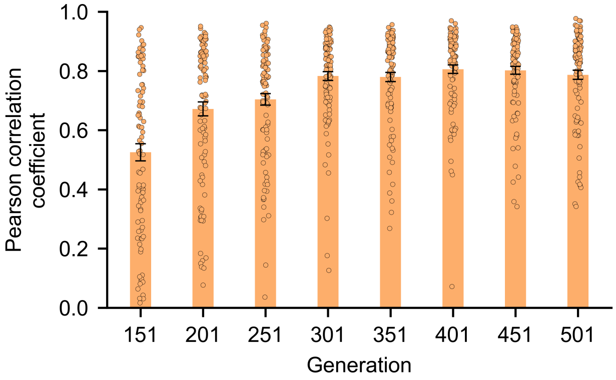


**Figure S9. LD inference via SLAFIT is accurate throughout the evolutionary history.** The later time-points exhibit stronger values of the Pearson correlation coefficient because these contain more variation which improves the accuracy of the haplotype reconstruction process. The bars show the mean Pearson correlation coefficient over 100 Monte Carlo runs. Each Monte Carlo run consisted of evolving populations of $N = 1\,000$ individuals of genome length $L = 4\,500$ bp. The genomes consisted of bi-allelic (WT and mutant) loci with equal forward and backward SNV probabilities set to $\mu = 10^{-4}$ per locus per generation. Alleles at $180/180/4140$ loci were respectively beneficial/deleterious/neutral with selection coefficients $+0.025/-0.025/0$. The beneficial and deleterious mutants were uniformly distributed across the entire genome. One founder sequence was used to generate each population, which was allowed to evolve for 550 generations and sampled at $\Delta t = 50$ generations. TGS reads of mean length 1 500 bp were simulated with 10% error rate and 50X coverage. The error bars show the standard error of the mean.

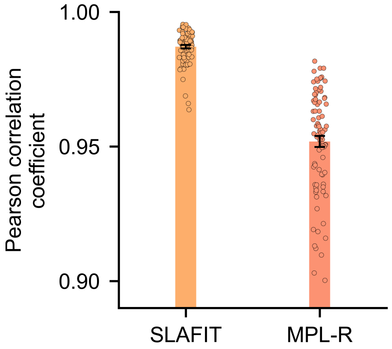


**Figure S10. The association between the estimates of SLAFIT and the ground truth values is stronger as compared to an NGS-based scheme.** The bars indicate the mean Person correlation coefficient values. The error bars indicate the standard error of the mean. The statistics are shown for 100 Monte Carlo runs. Each Monte Carlo run consisted of evolving populations of $N = 1\,000$ individuals of genome length $L = 4\,500$ bp. The genomes consisted of bi-allelic (WT and mutant) loci with equal forward and backward SNV probabilities set to $\mu = 10^{-4}$ per locus per generation. Alleles at $180/180/4140$ loci were respectively beneficial/deleterious/neutral with selection coefficients $+0.025/-0.025/0$. The beneficial and deleterious mutants were uniformly distributed across the entire genome. One founder sequence was used to generate each population, which was allowed to evolve for 550 generations and sampled at $\Delta t = 50$ generations. Unless otherwise stated, TGS reads of mean length 1 500 bp were simulated with 10% error rate and 50X coverage. 14 500 NGS reads of length 150 bp were simulated with to obtain similar coverage as the TGS reads.

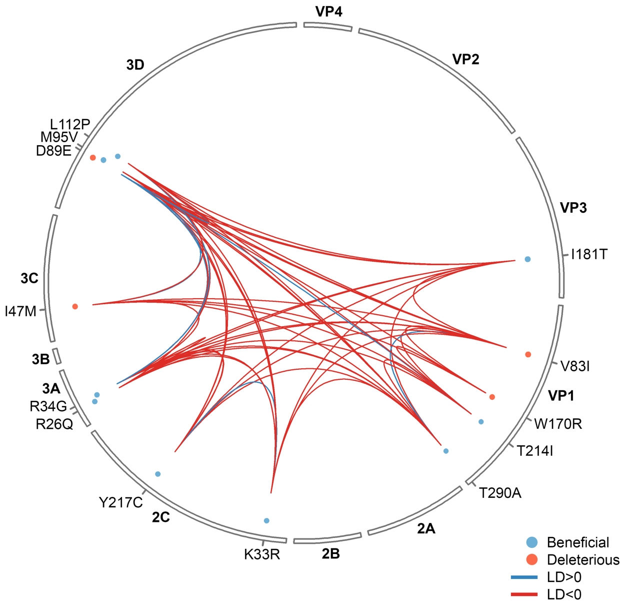


**Figure S11. LD patterns in the EV-C104 infection data show evidence of clonal interference.** log magnitudes of the inferred selection coefficients are indicated as the radii of the circles. Blue circles represent positive values, whereas red circles represent negative ones. Links between loci indicate LD values aggregated over all time-points. The widths of the link indicate the magnitude of the aggregated LD, and blue/red colors indicate positive/negative sign of the aggregated LD.

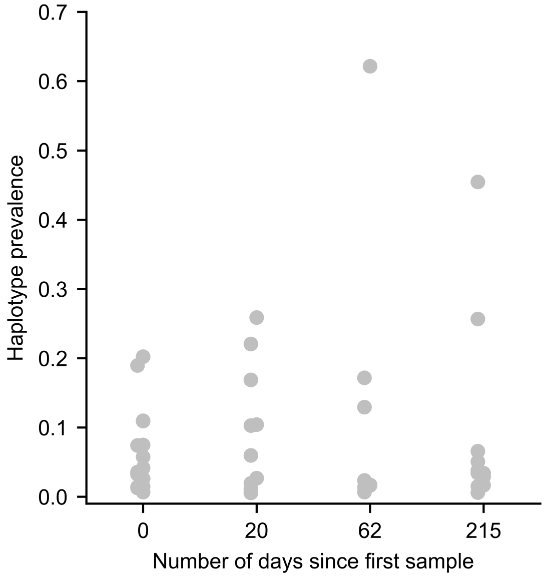


**Figure S12. Prevalence values of haplotypes reconstructed from the EV-C104 infection data indicate decreasing diversity with time.** The first two time-points show multiple haplotypes with similar prevalence values, whereas the last two time-points show a segregation between dominant haplotypes and the rest of the population.

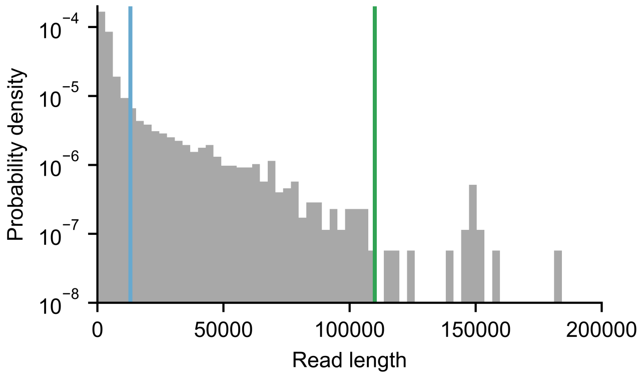


**Figure S13.** Distribution of read lengths in the HSV-1 dataset. Multiple reads are longer than the length of the $U_S$ region (blue line) and can capture SV in it. Some ultralong reads are longer than the length of the $U_L$ region (green line).
